# Algal turf sediment dynamics across the Florida Reef Tract

**DOI:** 10.64898/2026.11.02.763801

**Authors:** Manuel Ploner, Nicole L. Messerlian, Nathaniel Nutting-Hartman, Sterling B. Tebbett, Tory J. Chase, Fraser A. Januchowski-Hartley, Andrew G. Bauman

## Abstract

Algal turfs and their associated sediments are dominant components of many degraded reefs, where they mediate key ecosystem processes. However, the quantification of algal turfs and their sediments across reef shelves is limited outside Indo-Pacific reef systems. We quantified algal turf length, sediment loads, and sediment grain size distribution across four reef positions (nearshore, inner, middle, and outer) at three locations along the northern Florida Reef Tract (FRT). Sediment loads were positively associated with algal turf length and generally declined moving offshore. Nearshore reefs supported algal turfs that were both 1.6-fold longer and contained 1.6-fold higher sediment loads than on outer reefs, while sediment composition shifted toward finer grain sizes offshore. Nevertheless, regardless of reef position, all algal turfs had high sediment loads ranging from 1,445 to 16,639 g m⁻². Our findings suggest that cross-shelf patterns in algal turf sediment dynamics are present on Western Atlantic reefs and reveal that long sediment-laden algal turfs dominate on the FRT. These results highlight the pervasive nature of sediment-laden algal turfs on degraded reefs and emphasize the need to monitor and manage their expansion on reefs subject to chronic coastal development and climate-driven disturbance.

## 1. Introduction

Anthropogenic stressors are increasingly reshaping marine ecosystems (Hughes et al., 2017; Wernberg et al., 2016), often reducing topographic complexity through the loss of habitat-forming organisms and increased cover of low-lying taxa (Alvarez-Filip et al., 2009; O’Brien and Scheibling, 2018; Pessarrodona et al., 2021). Coral reefs exemplify this trend, with a long history of local stressors impacting reef communities, including overfishing (Cramer and Kittinger, 2021; Zamborain-Mason et al., 2023), pollution, local population growth and increased sediment inputs (Andrello et al., 2022; Arias-Gonzalez et al., 2017; Wenger et al., 2020). Further, over the past 40 years, increased frequency of repeated global coral bleaching and mass-mortality events have further driven reefs towards lower habitat complexity (Graham et al., 2006; Hughes et al., 2018; Reimer et al., 2024). Many reefs are now characterized by low coral cover (Acosta-Chaparro et al., 2025; Bellwood et al., 2019), with low structural complexity (Alvarez-Filip et al., 2009), and high algal turf cover (Tebbett et al., 2023a), which together increases sediment accumulation and retention, with likely negative consequences for reef ecosystem function (Tebbett et al., 2019).

Despite the increasing dominance of these modern reef habitats, the interactions between algal turfs and sediment, and the ecological roles of algal turfs remain understudied. Algal turfs are short (< 2 cm), multi-species assemblages of generally filamentous macroscopic algae that cover hard reef substrata and are broadly classified as short productive algal turfs (SPATs: < 3-5 mm in length with low volume of sediment) or long sediment-laden algal turfs (LSATs: > 3-5 mm in length with high volume of trapped sediment) (Goatley et al., 2016; Guzman et al., 2026; Tebbett et al., 2024). Algal turfs, particularly SPATs, drive benthic productivity on reefs (Carpenter, 1986; Klumpp and McKinnon, 1992), supporting key trophic pathways (Russ, 2003; Tebbett et al., 2018a; Wilson et al., 2003). However, increasing turf length enhances sediment retention, promoting a shift toward LSATs (Duran et al., 2024; Goatley et al., 2016), which impede coral settlement, development, and recovery (Birrell et al., 2005; Duran et al., 2024; Ford et al., 2018; Speare et al., 2019). Sediment-laden turfs are also associated with reduced grazing (Bellwood and Fulton, 2008; Goatley and Bellwood, 2012), resulting in a self-reinforcing feedback loop where increasing sediment loads promote turf growth, which further enhances sediment retention and further inhibits herbivory and coral recruitment (Birrell et al., 2005; Ladd et al., 2025; Syafruddin et al., 2025; Tebbett and Bellwood, 2019).

Importantly, turf-sediment interactions vary along environmental gradients across reef systems (Goatley and Bellwood, 2012; Guzman et al., 2026; Tebbett et al., 2018b). Research from the GBR (Tebbett et al., 2025; Tebbett et al., 2018b) and Indonesia (Syafruddin et al., 2025) has revealed that algal turf sediment dynamics vary along cross-shelf gradients. For example, nearshore reefs are typically characterized by high sediment loads and greater algal turf dominance, reflecting the influence of terrestrial inputs, high sediment retention, reef geomorphology, and often low hydrodynamic activity (Syafruddin et al., 2025; Tebbett et al., 2017; Tebbett et al., 2018b). In contrast, offshore reefs generally experience reduced terrestrial inputs and higher hydrodynamic energy which interacts with reef geomorphology to limit sediment accumulation associated with shorter algal turfs (Syafruddin et al., 2025; Tebbett et al., 2025; Tebbett et al., 2017). Given the links between algal turfs and key ecological processes on reefs, such as coral recruitment (Arnold et al., 2010; Birrell et al., 2005) and herbivory (Bellwood and Fulton, 2008; Goatley and Bellwood, 2012), such spatial variation in algal turf sediment dynamics may be reflected in bottom-up effects on reef functioning cross-shelf. However, our current understanding of cross-shelf algal turf sediment dynamics remains limited to Indo-Pacific reefs. Whether these patterns extend to distinct reef systems, such as those in the Western Atlantic, remains unclear, limiting broader understanding of reef functioning across biogeographic regions.

Compared to Indo-Pacific reefs, Western Atlantic reefs generally support lower biodiversity (McWilliam et al., 2018; Parravicini et al., 2013), reduced functional redundancy (Gahan et al., 2026; McWilliam et al., 2018), and lower resilience (Kuffner and Toth, 2016; Roff and Mumby, 2012), alongside distinct community composition and ecosystem functioning (Bellwood et al., 2004; Roff and Mumby, 2012). This suggests that Indo-Pacific ecological patterns may not extend to the Western Atlantic, underscoring the need for region-specific investigations of turf– sediment dynamics.

The Florida Reef Tract (FRT) supports a pronounced cross-shelf gradient from nearshore to offshore reefs with spatial variation in benthic community composition, environmental conditions, and hydrodynamic regimes (Banks et al., 2008; Moyer et al., 2003). Florida reefs are now characterized by extensive algal turf cover, high sediment loads, and low structural complexity (Alvarez-Filip et al., 2009; Guzman et al., 2026), which are expected to promote LSAT development and persistence (Ladd et al., 2025). Examining these dynamics in this high-latitude, marginal reef system would extend understanding beyond the Indo-Pacific and help assess the generality of cross-shelf patterns in the Western Atlantic. Consequently, examining these dynamics on the FRT would not only extend our understanding beyond the Indo-Pacific, but also provide insights into cross-shelf turf–sediment dynamics in a high-latitude reef system. Such insights are increasingly relevant as ocean warming promotes the expansion of turf-dominated states on marginal and temperate reefs (Agostini et al., 2021; Vergés et al., 2019).

This study characterized spatial variation in algal turf length and associated sediment loads across the FRT. We quantified turf length, sediment loads, and sediment grain size distributions on adjacent nearshore, inner, middle, and outer reefs at three locations to examine cross-shelf patterns and potential links between sediment characteristics and LSAT development. Collectively, these analyses provide the first characterization of cross-shelf turf–sediment dynamics in a high-latitude Western Atlantic reef system and provide a foundation for investigating how turf–sediment interactions shape reef degradation and resilience.

## 2. Materials and methods

### 2.1. Study sites

This study was conducted between October 2024 and March 2025 in the northern portion of the FRT, United States of America. The study locations Pompano Beach (26.2363, -80.0820), Lauderdale Beach (26.1569, -80.0957), and Hollywood (26.0079, -80.1086), were selected because they are approximately equidistant along a broad stretch (∼25 km) of the northern FRT. At each location, four reef positions were sampled including the nearshore, inner, middle, and outer reef, with one site per position (n = 12 sites, Fig. 1). Reefs were characterized by relatively low structural complexity and were visually dominated by algal turfs. Depths and distances to shore varied predictably across the shelf gradient, with nearshore reefs occurring at approximately 4–7 m depth and 0.6 km from shore, inner reefs at 7–13 m and 1.1 km, middle reefs at 12–16 m and 1.7 km, and outer reefs at 16–20 m and 2.3 km. Depth and distance from shore covary naturally across the FRT; our sampling design therefore characterized spatial patterns in turf-associated sediments without isolating the effects of these factors.

**Fig. 1.**
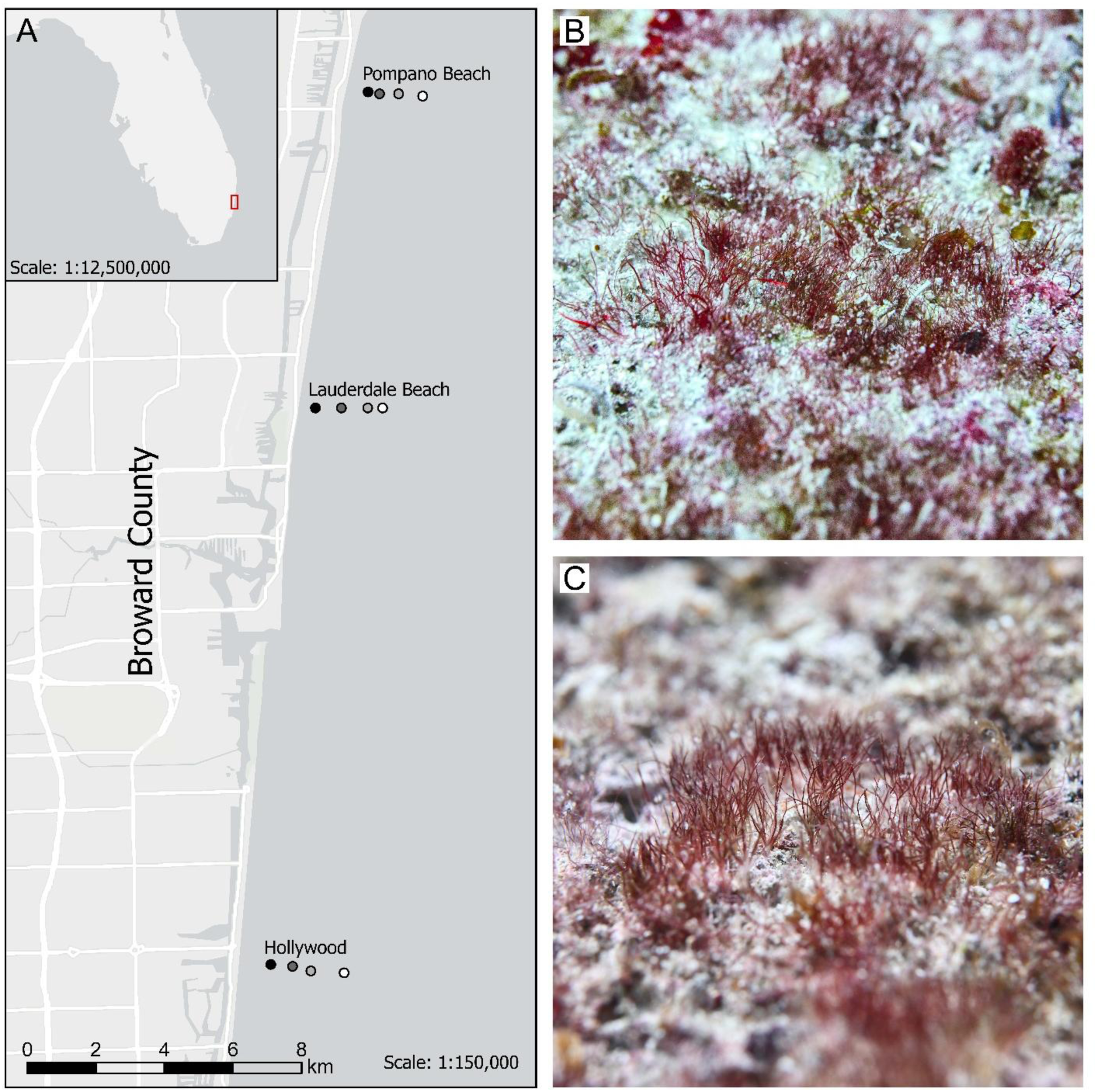
Study area and representative algal turf communities along the FRT. (**A**) Map of the study region in Broward County, Florida, USA, showing the three sampling locations (Pompano Beach, Lauderdale Beach, and Hollywood) and four reef positions sampled at each location (nearshore – black, inner reef – dark grey, middle reef – light grey, and outer reef – white) on the FRT. (**B-C**) Algal turf community following sediment removal at Lauderdale Beach outer reef **(B**) and inner reef site (**C**).

### 2.2. Sediment sampling & algal turf measurement

Particulate material (sediments and associated organic material) within algal turfs was collected using an underwater vacuum from within a 58 cm² PVC ring placed on a suitable algal turf covered surface following Tebbett et al. (2022). Suitable surfaces were defined as flat, mostly non-porous areas of hard reef substratum located outside territories of farming damselfishes and free of sediment-retaining pits, macroalgae, and encrusting organisms. At each site, 10 sediment samples were collected (n = 120 samples across sites), with replicates separated by a minimum of five meters. Following sediment removal, turf height was measured for 10 haphazardly selected algal turf filaments within each 58 cm^2^ PVC sampling area using the depth probe of vernier calipers.

### 2.3. Sediment processing

Individual sediment samples were transferred to 9 L containers, and 15 mL of 10% buffered formalin was added to fix organic material. Samples were then allowed to settle for at least 5 h to ensure particulate material >10 µm had settled (Purcell, 1996), after which the overlying water was decanted and the remaining sediment transferred to a 473 mL container. An additional 10 mL of buffered formalin was added to the 473 ml container to preserve samples during the extended processing. After another settling period of at least 5 h, the overlying water was decanted, and the sediments were transferred to a 175 mL sample jar containing 5 mL of formalin. All samples were then wet sieved through a 2000 µm plastic mesh to remove coarse particulate material >2 mm, retaining the sand, silt, and clay fractions (ISO 14688-1:2017). Samples were then rinsed three times with deionized water to remove residual salts and prevent salt crystal formation, allowing at least 5 h of settling between rinses.

Following the final rinse, samples were dried to a constant weight at 60°C and weighed to the nearest 0.0001 g to determine total particulate mass. Organic material was then removed using trace-metal-grade 35% hydrogen peroxide (H_2_O_2_) for a minimum of two weeks, with samples stirred and replenished with 10–15 ml of fresh H_2_O_2_ every two days to ensure complete oxidation. Samples were subsequently re-dried and re-weighed to determine total sediment mass and were standardized to g m⁻². Sediment grain-size fractionation was then conducted using plastic mesh sieves to separate the following size classes: <63 µm, 63–125 µm, 125–250 µm, 250–500 µm, 500–1000 µm, and 1000–2000 µm, which were individually weighed.

### 2.4. Statistical analysis

Statistical analyses were conducted in R (version 4.5.1; R Core R-Core-Team, 2025) for univariate and mixed-effects modeling, and in PRIMER v7 with the PERMANOVA+ add-on (Clarke and Gorley, 2015) for multivariate analyses. To examine spatial patterns in algal turf length and sediment mass, and their relationship across Florida’s shelf, we fitted generalized linear mixed models (GLMMs) with a Gamma distribution and log-link function using the *glmmTMB* package (Brooks et al., 2017). Model validation was performed using the *DHARMa* package (Hartig, 2024), and marginal and conditional *R²* values were calculated using the *performance* package (Lüdecke et al., 2021). Estimated marginal means (EMMs) were obtained on the response scale using the *emmeans* package (Lenth and Piaskowski, 2025), and Tukey-adjusted pairwise comparisons were used to compare between factor levels.

First, we examined spatial variation of algal turf length and sediment mass, independently from each other, as a function of reef position (nearshore, inner, middle and outer reefs) and location (Pompano Beach, Lauderdale Beach and Hollywood). Second, to examine the relationship between algal turf length and sediment load and explore variation in the length of algal turfs per unit of sediment mass, we modeled algal turf length as a function of log-transformed sediment mass, with reef position and location as fixed effects. All models included site as a random effect to account for spatial non-independence among samples collected within sites. Interaction terms were evaluated during model selection but were excluded from final models because models including interaction consistently resulted in higher AICc values and/or showed signs of overparameterization. Model fit was assessed using simulated quantile residuals, including residual-versus-fitted plots and residual patterns across reef position, location, sediment mass, and site. Residual diagnostics indicated acceptable model fit for all models.

Sediment grain-size composition was expressed as the proportional contribution of the six grain-size fractions to total sediment mass and analyzed as a multivariate dataset. Differences in sediment composition among reef positions and locations were tested using permutational multivariate analysis of variance (PERMANOVA) on a Euclidean distance matrix of the proportional data. The PERMANOVA included reef position and location as fixed factors, with site treated as a random effect nested within the position × location interaction. Type III sums of squares and 9,999 permutations under a reduced model were used. Because the reef position × location interaction was significant, pairwise tests were used to compare reef positions separately within each location. Homogeneity of multivariate dispersion was assessed using PERMDISP with 9,999 permutations, both across the full dataset and separately within each location to evaluate potential differences in dispersion among reef positions.

Principal component analysis (PCA) was used to visualize multivariate variation in sediment composition among reef positions and identify the grain-size fractions contributing most strongly to cross-shelf patterns. Further, similarity percentage analysis (SIMPER) was conducted using Bray–Curtis similarity matrices calculated from raw proportional data. SIMPER was used to quantify the relative contribution of each grain-size fraction to dissimilarities among reef positions and locations.

## 3. Results

### 3.1. Spatial patterns in algal turf length

Turf length ranged from 2.6 to 14.6 mm (mean = 6.40 ± 0.21 mm) and exhibited clear spatial variation across the reef system (Fig. 2 A and B). Algal turfs were significantly shorter on outer reefs (4.76 ± 0.35 mm) than the middle reef (6.43 ± 0.47 mm, *p* < 0.02), inner reef (6.22 ± 0.45 mm, *p* < 0.05), and nearshore positions (7.71 ± 0.56 mm, *p* < 0.0001). No differences were detected between nearshore, inner reef, and middle reef positions (all *p* > 0.15; Supplementary Table S1). Alongshore variation was also evident: turfs were 1.52-fold longer at Pompano Beach and 1.26-fold longer at Lauderdale Beach than at Hollywood (p < 0.0001 and p < 0.03, respectively). Turf length did not differ significantly between Pompano Beach and Lauderdale Beach (p > 0.05; Supplementary Table S1). Overall, the additive GLMM explained a substantial proportion of variation in turf length (marginal R² = 0.49; conditional R² = 0.58).

**Fig. 2.**
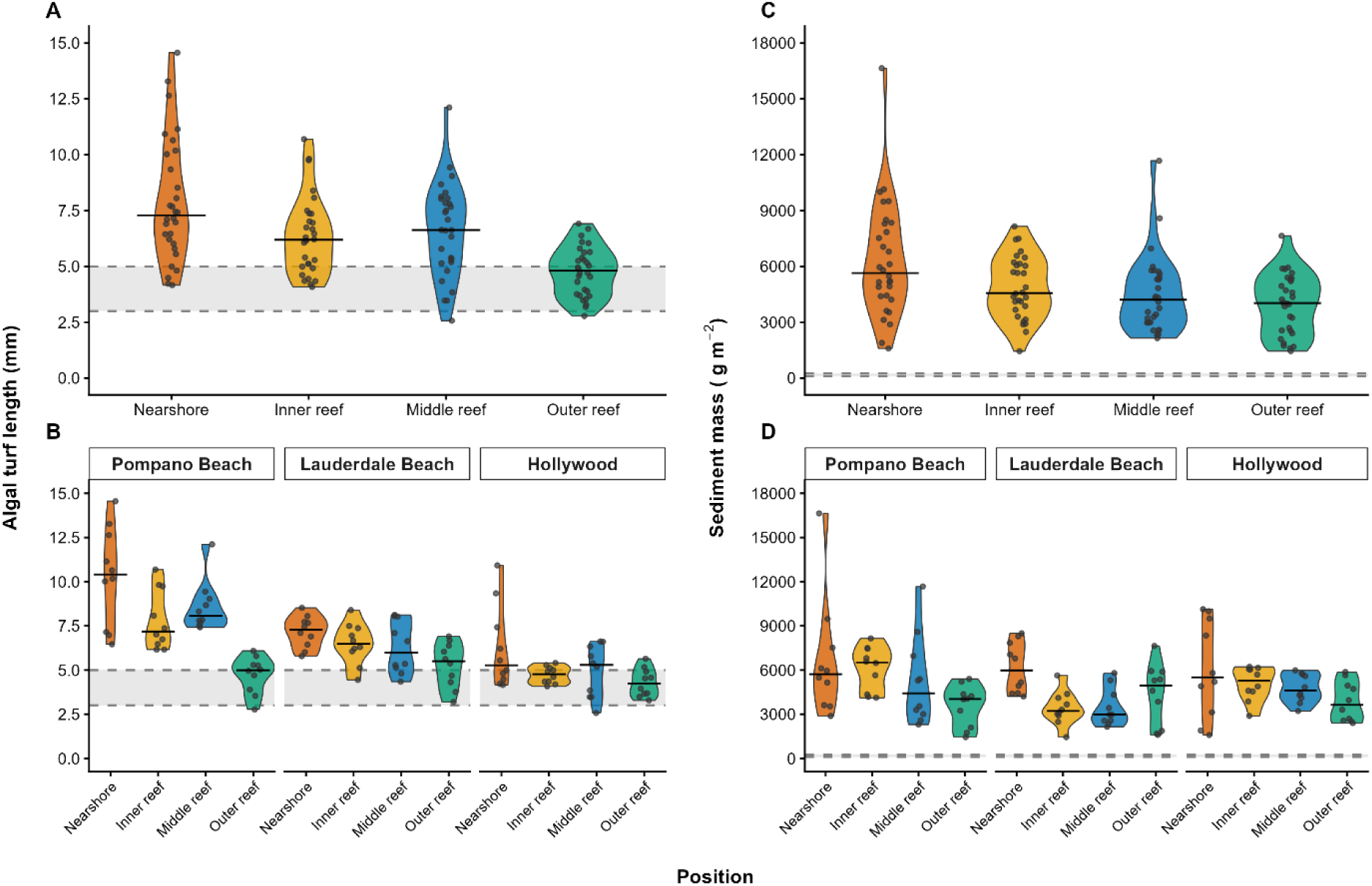
Spatial variation on the FRT for algal turf length (mm) and sediment mass (g m⁻²) across reef position (**A, C**) and location (**B, D**). Dashed horizontal lines and shaded areas indicate the 3-5 mm (**A**, **B**) or the 100-250 g m⁻² threshold (**C**, **D**) used to define LSATs. In all graphs, violin plots show the data distributions, horizontal bars indicate medians, and points represent individual samples. Colors denote reef positions: nearshore (orange), inner reef (yellow), middle reef (blue), and outer reef (green).

### 3.2. Spatial patterns in sediment mass

Sediment mass ranged from 1,445 to 16,639 g m⁻² (mean = 4,896 ± 212 g m⁻²) across all samples and declined with increasing distance from the shore (Fig. 2 C and D). Nearshore reefs (6,281 ± 663 g m⁻²) had significantly greater sediment loads/mass than outer reefs (3,939 ± 417 g m⁻², *p* < 0.01). No other pairwise differences were detected between nearshore, inner reef (4,750 ± 502 g m⁻²), middle reef (4,424 ± 467 g m⁻²), and outer reef positions (all *p* > 0.08; Supplementary Table S2). Sediment mass did not differ between locations (all pairwise comparisons *p* > 0.20; Supplementary Table S2). Overall, reef position and location explain a modest proportion of the variation in sediment mass (marginal R² = 0.18; conditional R² = 0.26).

### 3.3. Algal length co-variation with sediment mass

Algal turf length was positively associated with sediment mass (β = 0.229 ± 0.046 SE, z = 4.98, p < 0.001; Fig. 3). Differences among reef positions remained significant after accounting for sediment mass, and the model explained 57% of the variation through fixed effects and 65% when the random effect of site was included (marginal R² = 0.57; conditional R² = 0.65). Predicted algal turf length was 1.45 times greater on nearshore and 1.30 times greater on middle reefs compared to outer reefs (p < 0.002 and p < 0.04, respectively). No significant differences were detected among nearshore, inner, and middle reef positions or between inner and outer reefs (all p > 0.12; Supplementary Table S3).

**Fig. 3.**
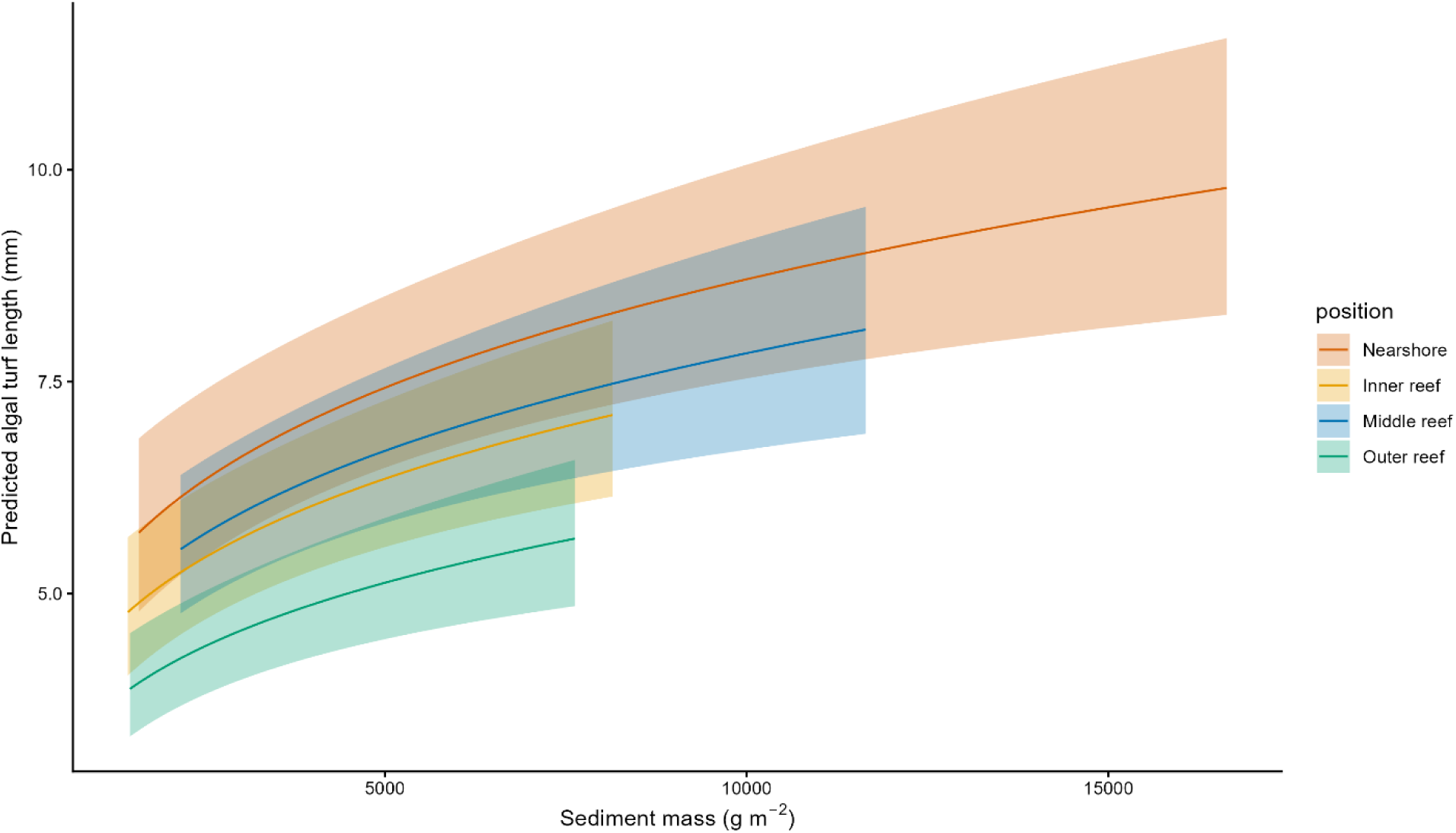
Model predicted relationship between sediment mass (g m⁻²) and algal turf length (mm) across reef positions on the Florida Reef Tract based on a generalized linear mixed model. Colored lines represent predicted mean turf length for each reef position and shaded ribbons indicate 95% confidence intervals.

Alongshore patterns persisted after accounting for sediment mass. Predicted algal turf length was 1.51 times greater at Pompano Beach and 1.31 times greater at Lauderdale Beach compared to Hollywood (p < 0.0001 and p < 0.005, respectively). No significant difference was detected between Pompano Beach and Lauderdale Beach (p > 0.19; Supplementary Fig. S1. and Table S3).

### 3.4. Spatial patterns in sediment size fractionation

Sediment grain-size composition varied significantly across the reef system (Fig. 4), with differences among reef positions (PERMANOVA, pseudo-F₃,₁₀₈ = 58.18, *p* < 0.0001), locations (pseudo-F₂,₁₀₈ = 32.74, *p* < 0.0001), and their interaction (pseudo-F₆,₁₀₈ = 15.39, *p* < 0.0001; Supplementary Table S4). Pairwise comparisons revealed significant differences among all reef positions within Pompano, Lauderdale Beach, and Hollywood (all *p* < 0.002), although compositional differences between middle and outer reefs were less pronounced than nearshore and offshore reefs (Supplementary Fig. S2. and Table S5).

**Fig. 4.**
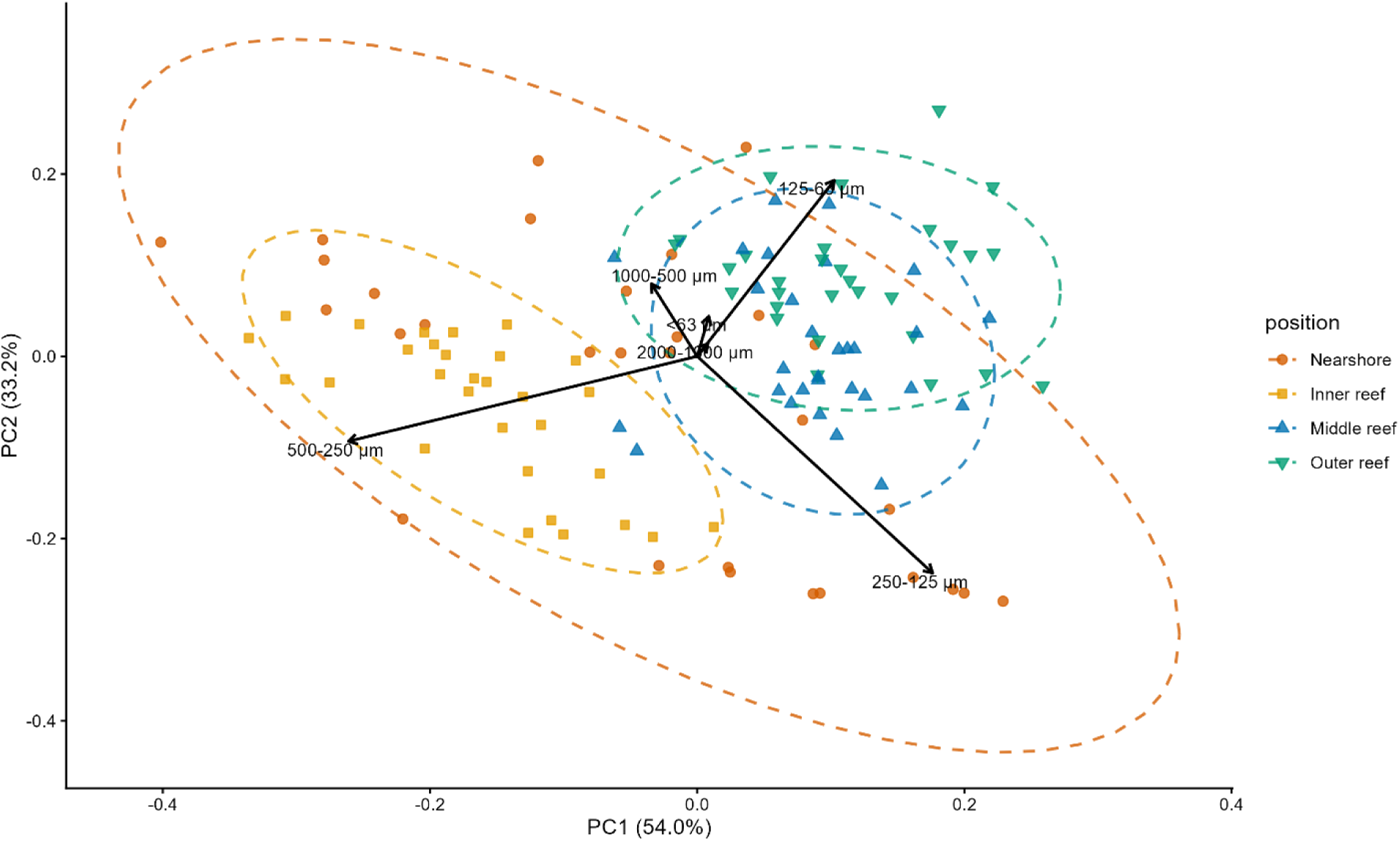
Principal component analysis (PCA) of proportional sediment grain-size composition across reef positions on the Florida Reef Tract. The first two principal components explained 87.2% of the total variation in sediment composition (PC1 = 54.0%, PC2 = 33.2%). Points represent individual sediment samples and dashed ellipses indicate group dispersion by reef position: Arrows represent eigenvectors of sediment grain-size fractions, illustrating the direction and relative contribution of each fraction to multivariate variation in sediment composition.

Multivariate dispersion differed significantly among reef positions (PERMDISP, F₃,₁₁₆ = 41.19, *p* < 0.0001), with nearshore reefs exhibiting the greatest variability in sediment grain-size composition (Fig. 4) and therefore, the main effect of reef position was interpreted cautiously. However, dispersion did not differ significantly among locations (F₂,₁₁₇ = 0.33, *p* > 0.73) or among reef positions within individual locations (all *p* > 0.06; Supplementary Table S6).

SIMPER analysis indicated that cross-shelf differences in sediment composition were driven primarily by shifts in fine and intermediate sediment grain-size fractions (Fig. 4). The 63–125 µm size class consistently contributed strongly to dissimilarities involving offshore reefs, particularly outer reefs, whereas the 250–500 µm fraction was an important driver of contrasts involving nearshore reefs (Fig. 4). The 500–1000 µm and 125–250 µm fractions also contributed consistently to cross-shelf and alongshore dissimilarities, while the coarsest and finest fractions (< 63 µm; 1000–2000 µm) contributed comparatively little. Overall, nearshore and inner reefs contained higher proportions of intermediate grain-sized sediments, while middle and outer reefs contained higher proportions of finer sediments.

## 4. Discussion

Our study revealed cross-shelf differences in algal turf length and sediment mass along the Florida Reef Tract (FRT), with both generally declining from nearshore to outer reefs. Moreover, algal turf length was also strongly and positively associated with sediment mass, while sediment composition shifted systematically toward finer grain sizes offshore. Despite the much smaller cross shelf extent of the FRT (nearshore to offshore reef distance of 2 – 3 km compared to > 30 for both the Spermonde Archipelago and Great Barrier Reef (GBR)), the cross-shelf patterns we found here are similar in kind, although sediment loads vary substantially between regions (Syafruddin et al., 2025; Tebbett et al., 2025; Tebbett et al., 2018b). Specifically, mean sediment load on nearshore FRT reefs was, at almost 6.3 kg m^-2^, approximately three times greater than sediment loads reported from coastal reefs on the GBR (Tebbett et al., 2018b), and more than eighty times greater than the mean reported from the Spermonde Archipelago (Syafruddin et al., 2025). Despite these exceptionally high sediment loads, measured algal turf lengths on FRT nearshore reefs (7.89 ± 0.50 mm) were comparable to lengths reported from other sediment-laden reef environments including the GBR (Goatley et al., 2016), the Dampier Archipelago (Tebbett et al., 2024), and Puerto Rico (Tebbett et al., 2026). While the positive association between turf length and sediment load appears consistent across reef systems (e.g. Goatley et al., 2016; Syafruddin et al., 2025; Tebbett et al., 2024), sediment retention within algal turfs varies substantially among regions, potentially reflecting differences in environmental conditions, reef geomorphology, and local disturbance regimes.

Although the sources of sediments measured in this study could not be determined directly, the observed offshore decline in sediment loads, together with evidence from other nearby ports such as Miami (Miller et al., 2016), suggest that terrestrial and dredging-derived inputs may contribute to the elevated sediment loads. Southeast Florida’s coastline is heavily urbanized, and chronic local stressors (e.g., coastal construction, dredging, port operations, beach nourishment) drive sustained sediment inputs (Enochs et al., 2019; Jordan et al., 2010; Miller et al., 2016; Rogers and Ramos-Scharrón, 2022). For example, at Port Miami, sediment cover on consolidated hardbottom habitat at inner-reef sites affected by dredging at Port Miami was 17.5– 36 times greater than at reference sites (Miller et al., 2016), and extended at least 2.5 km from the channel and persisted for several years (Serrano et al., 2026). Together with natural cross-shelf sediment transport and retention processes, these inputs may explain the high sediment loads on the northern FRT and their decline with increasing distance from shore.

Cross-shelf variation in light, hydrodynamic energy, and reef geomorphology, may also shape algal turf growth and sediment dynamics (Mueller et al., 2016; Schlaefer et al., 2021; Tebbett et al., 2018b; Torres-Garcia et al., 2018). Higher irradiance on shallow nearshore reefs can promote algal turf growth, while wave-driven sediment resuspension can sort sediments and preferentially remove finer sediments (Storlazzi et al., 2004; Tebbett et al., 2018a; Torres-Garcia et al., 2018; Wolanski et al., 2005). Conversely, reduced light and lower near-bed hydrodynamic energy on deeper outer reefs may constrain algal turf growth and facilitate the deposition and retention of finer sediments within the turf matrix (Mueller et al., 2016; Tebbett et al., 2018b; Torres-Garcia et al., 2018). For example, Reef geomorphology may further modify these patterns: low-relief nearshore reef habitats are more exposed to sediment transported from adjacent unconsolidated substrates, whereas the greater relief and structural complexity of offshore reefs can alter local transport, deposition, and retention (Schlaefer et al., 2021; Schlaefer et al., 2022; Tebbett et al., 2023b). These mechanisms are consistent with our findings that sediment loads generally declined offshore, while sediment composition shifted from intermediate grain sizes (250–500 µm) on nearshore and inner reefs toward finer sediments (63–125 µm) on outer reefs.

While sediment mass remained relatively consistent between locations, algal turf length and sediment composition varied, yet general cross-shelf patterns in algal turf length and sediment composition persisted. Algal turfs at Pompano Beach were ∼50% longer than those at Hollywood despite similar sediment loads, suggesting that factors beyond sediment quantity influence local algal turf development. Alongshore variation may reflect local differences in hydrodynamic conditions (Storlazzi et al., 2004; Williams and Carpenter, 1998), nutrient inputs (Benkwitt et al., 2025; Fong et al., 2018), and proximity to coastal infrastructure (e.g., inlets and ports; Enochs et al., 2019), which can affect sediment transport and nutrient availability. Thus, local conditions appear to modulate sediment-algal turf dynamics, without obscuring broader cross-shelf gradients across the northern FRT.

Given that algal turf-associated sediments can shape multiple reef processes (Adam et al., 2018; Ladd et al., 2025; Storlazzi et al., 2015), the high sediment loads documented across the FRT, particularly on nearshore reefs, may have broad consequences for ecosystem functioning. Sediment accumulation can directly alter algal turf communities (Clausing et al., 2014; Fong et al., 2018): as turfs stabilize sediment within their matrix, longer filaments may trap additional sediment, organic material, and nutrients, promoting LSAT development through a self-reinforcing feedback (Guzman et al., 2026; Ladd et al., 2025; Tebbett and Bellwood, 2019). The strong positive association between algal turf length and sediment mass observed in this study is consistent with this proposed feedback. LSATs persistence may be further reinforced through reduction in herbivore pressure, as herbivorous fishes graze sediment-laden turfs less intensively (Goatley and Bellwood, 2012; Ladd et al., 2025; Ng et al., 2021). This may be due to sediment accumulation reducing algal turf productivity and nutritional quality (Tebbett et al., 2018a), limiting the palatability and value of LSATs to herbivorous fishes (Benkwitt et al., 2025; Tebbett et al., 2024). Furthermore, accumulated sediments can impair coral recruitment and recovery (Birrell et al., 2005; Duran et al., 2024; Ford et al., 2018; Speare et al., 2019). The prevalence of LSATs and exceptionally high sediment loads across the northern FRT may therefore constrain both natural coral recovery and restoration success, particularly when restoration efforts do not address the conditions promoting sediment accumulation and algal turf persistence. Collectively, these interacting processes may perpetuate degraded reef states and diminish ecosystem functional processes and resilience associated with coral dominated reefs.

Our findings provide further evidence that current classifications distinguishing short productive algal turfs (SPATs) from long sediment-laden algal turfs (LSATs) may not adequately characterize turf communities on the FRT, where sediment loads greatly exceeded conventional LSAT thresholds even when turf lengths remained within or near the SPAT range. Outer-reef turfs, for example, averaged <5 mm in length but contained approximately 4,000 g m⁻² of sediment, revealing a clear mismatch between turf structure and sediment accumulation under the existing framework (see also Fig. 6a in Tebbett et al., 2024). Similarly, recent research from the FRT suggests that the transition between SPATs and LSATs may occur at shorter turf lengths and sediment depths of 2–3 mm (Guzman et al., 2026). Although the SPAT–LSAT framework has proven useful on reef systems such as the Great Barrier Reef, additional categories may be needed to capture context-dependent algal turf states. These could include short sediment-laden algal turfs (SSATs) and long productive algal turfs (LPATs), such as those found within damselfish territories (Klumpp et al., 1987; Tebbett et al., 2020). Expanding the existing framework to incorporate such context-dependent turf states may improve our ability to compare algal turf dynamics across reef systems and better capture the functional diversity of turf communities under varying environmental conditions.

Although our study identified strong spatial patterns in algal turf length, sediment load, and sediment composition across the northern Florida Reef Tract, several limitations should be acknowledged. Sampling captured a single point in time and therefore did not account for temporal variability, episodic sedimentation events, or storm-driven disturbance that may influence reef sediment dynamics and benthic communities (Cecchetto et al., 2024; Scoffin, 1993; Wolanski et al., 2008). While quantification of temporal variation in turf-sediment dynamics warrants future investigation, initial research on this topic suggests that such variation could be relatively limited (Gordon et al., 2016). Moreover, several potential drivers of the observed cross-shelf patterns including reef geomorphology, hydrodynamic energy, light availability, and herbivory were not measured. Future studies should combine repeated sampling with measurements of grazing, water flow, sediment resuspension and deposition, nutrient availability, and benthic productivity. In this respect, experimental manipulations of sediment addition and removal, and herbivore exclusion could further determine how sediment retention and sediment-associated constituents (e.g., organic matter, nutrients and contaminants) influence algal turf development, coral recruitment, and herbivory. Together, these approaches would help clarify the mechanisms underlying algal turf sediment feedbacks and their consequences for reef functioning in increasingly urbanized coastal environments.

In summary, our findings reveal pronounced cross-shelf variation in algal turf length, sediment load, and sediment composition across the FRT, extending previous research for the western Atlantic. Exceptionally high sediment loads across the FRT, even on deeper offshore reefs, suggest that sediment-laden turf states may represent an important, but often overlooked, feature of marginal, urbanized reef systems. Because LSATs can reduce herbivory, suppress coral recruitment, alter benthic productivity, and reinforce degraded reef states (Birrell et al., 2005; Duran et al., 2024; Ford et al., 2018; Ladd et al., 2025; Speare et al., 2019), their expansion and persistence may have important consequences for reef resilience under chronic coastal development and climate-driven disturbance. Our results also suggest that existing classification frameworks may not adequately capture the range of turf-sediment states across reef systems, especially in cases where short turfs retain high sediment loads. Incorporating additional algal turf states, such as SSATs and LPATs, may improve comparisons among reef systems and predictions of ecosystem responses to environmental changes. Overall, these findings highlight the prevalence of sediment-laden algal turfs on degraded reefs and emphasize the importance of incorporating algal turf community structure and associated sediment loads into reef monitoring and management.

## Acknowledgments

We would like to thank Sofi El-Rass, Madi Androne, Emma Brennan, Patrick Idank, Bernhard Riegl Jr., Grace Triplett, Naya Mondrosch, Sienna Lazar, the crew from the REEFS Research Group at Nova Southeastern University, and the Coast to Coral Reef Environmental Dynamics Lab at Skidmore College for assisting with field and lab work for this project. Funding was provided by the Florida Department of Environmental Protection (AGB and FAJH), the National Coral Reef Institute (AGB and FAJH), Nova Southeastern University’s President’s Faculty Research and Development Grant (AGB), and the Gale Foundation (MP). All research activities were conducted under the Florida Fish and Wildlife Special Activity License (SAL-24-2665-SR). National Coral Reef Institute at Nova Southeastern University through the National Oceanographic and Atmospheric Administration (NA23NOS4690272). The views, statements, findings, conclusions, and recommendations expressed herein are those of the author(s) and do not necessarily reflect the views of NOAA.

## CRediT authorship contribution statement

**Manuel Ploner**: Writing – review & editing, Writing – original draft, Visualization, Methodology, Investigation, Funding acquisition, Formal analysis, Data curation, Conceptualization. **Nicole L. Messerlian**: Writing – review & editing, Methodology, Investigation. **Nathaniel Nutting-Hartman**: Writing – review & editing, Methodology, Investigation, Data curation. **Sterling B. Tebbett**: Writing – review & editing, Methodology, Formal analysis, Conceptualization. **Tory J. Chase**: Writing – review & editing, Resources, Methodology, Investigation, Data curation, Funding acquisition, Conceptualization. **Fraser A. Januchowski-Hartley**: Writing – review & editing, Resources, Methodology, Investigation, Data curation, Funding acquisition, Conceptualization. **Andrew G. Bauman**: Writing – review & editing, Resources, Methodology, Investigation, Data curation, Funding acquisition, Conceptualization.

## Appendix A. Supplementary data

Supplementary data to this article can be found online at: LINK

## Data availability

Raw data from detailed algal turf and sediment quantification has been uploaded as a supplementary excel file: https://doi.org/10.5281/zenodo.23771106. All other data are available from the corresponding author on reasonable request.

