## Supplemental Material for "Algal turf sediment dynamics across the Florida Reef Tract"

### 1. Figures

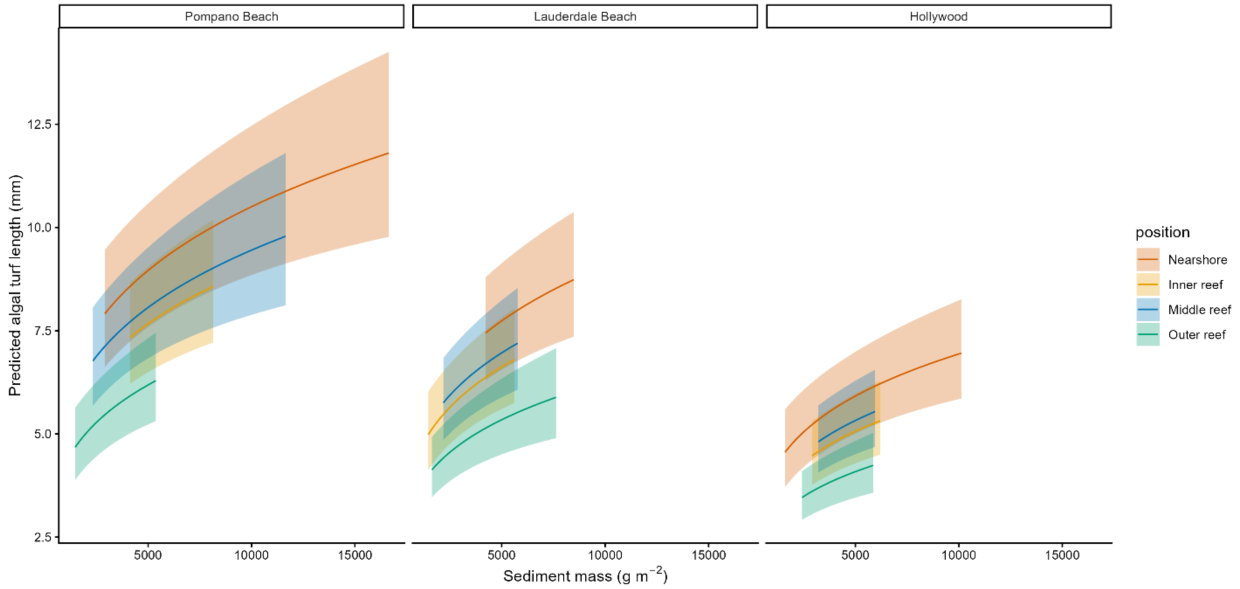


**Fig. S1.** Model predicted relationship between sediment mass (g m$^{-2}$) and algal turf length (mm) across reef positions at Pompano Beach, Lauderdale Beach, and Hollywood along the Florida Reef Tract based on a generalized linear mixed model. Colored lines represent predicted mean turf length for each reef position and shaded ribbons indicate 95% confidence intervals.

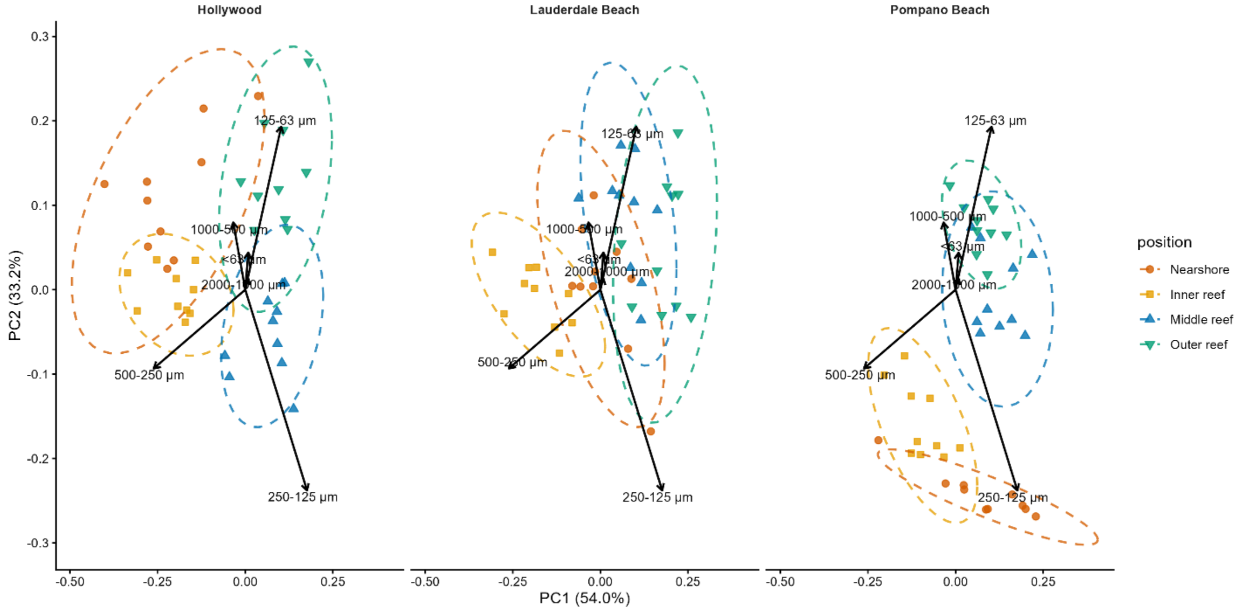


**Fig. S2.** Principal component analysis (PCA) of proportional sediment grain-size composition across reef positions at Hollywood, Lauderdale Beach, and Pompano Beach along the Florida Reef Tract. The first two principal components explained 87.2% of the total variation in sediment composition (PC1 = 54.0%, PC2 = 33.2%). Points represent individual sediment samples and dashed ellipses indicate group dispersion by reef position. Arrows represent eigenvectors of sediment grain-size fractions, illustrating the direction and relative contribution of each fraction to multivariate variation in sediment composition.

#### 2. Tables

Table S1. Pairwise comparisons of algal turf length among reef positions and locations based on estimated marginal means from a Gamma generalized linear mixed model (GLMM) with a log link function.

| Group | Comparison | Ratio | SE | z_ratio | *p*_adjusted |
|---|---|---|---|---|---|
| Position | Nearshore - Inner reef | 1.241 | 0.128 | 2.096 | 0.1545 |
| Position | Nearshore - Middle reef | 1.200 | 0.123 | 1.773 | 0.2864 |
| Position | Nearshore - Outer reef | 1.621 | 0.167 | 4.691 | <0.0001 |
| Position | Inner reef - Middle reef | 0.967 | 0.100 | -0.323 | 0.9884 |
| Position | Inner reef - Outer reef | 1.306 | 0.134 | 2.596 | 0.0465 |
| Position | Middle reef - Outer reef | 1.350 | 0.139 | 2.919 | 0.0185 |
| Location | Pompano Beach - Lauderdale Beach | 1.206 | 0.107 | 2.101 | 0.0896 |
| Location | Pompano Beach - Hollywood | 1.516 | 0.135 | 4.671 | <0.0001 |
| Location | Lauderdale Beach - Hollywood | 1.257 | 0.112 | 2.570 | 0.0275 |

Table S2. Pairwise comparisons of sediment mass among reef positions and locations based on estimated marginal means from a Gamma generalized linear mixed model (GLMM) with a log link function.

| Group | Comparison | Ratio | SE | z_ratio | *p*_adjusted |
|---|---|---|---|---|---|
| Position | Nearshore - Inner reef | 1.322 | 0.198 | 1.869 | 0.2414 |
| Position | Nearshore - Middle reef | 1.420 | 0.212 | 2.347 | 0.0877 |
| Position | Nearshore - Outer reef | 1.595 | 0.238 | 3.123 | 0.0097 |
| Position | Inner reef - Middle reef | 1.074 | 0.160 | 0.477 | 0.9643 |
| Position | Inner reef - Outer reef | 1.206 | 0.180 | 1.254 | 0.5926 |
| Position | Middle reef - Outer reef | 1.123 | 0.168 | 0.778 | 0.8646 |
| Location | Pompano Beach - Lauderdale Beach | 1.245 | 0.161 | 1.693 | 0.2079 |
| Location | Pompano Beach - Hollywood | 1.080 | 0.140 | 0.594 | 0.8233 |
| Location | Lauderdale Beach - Hollywood | 0.867 | 0.112 | -1.099 | 0.5145 |

Table S3. Pairwise comparisons of algal turf length among reef positions and locations after accounting for sediment mass, based on estimated marginal means from a Gamma generalized linear mixed model (GLMM) with a log link function.

| Group | Comparison | Ratio | SE | z_ratio | *p*_adjusted |
|---|---|---|---|---|---|
| Position | Nearshore - Inner reef | 1.169 | 0.115 | 1.589 | 0.3847 |
| Position | Nearshore - Middle reef | 1.111 | 0.110 | 1.069 | 0.7083 |
| Position | Nearshore - Outer reef | 1.449 | 0.145 | 3.701 | 0.0012 |
| Position | Inner reef - Middle reef | 0.951 | 0.093 | -0.518 | 0.9549 |
| Position | Inner reef - Outer reef | 1.239 | 0.122 | 2.183 | 0.1278 |
| Position | Middle reef - Outer reef | 1.303 | 0.128 | 2.708 | 0.0342 |

| | | | | | |
|---|---|---|---|---|---|
| Location | Pompano Beach - Lauderdale Beach | 1.158 | 0.098 | 1.726 | 0.1956 |
| Location | Pompano Beach - Hollywood | 1.514 | 0.128 | 4.911 | <0.0001 |
| Location | Lauderdale Beach - Hollywood | 1.308 | 0.111 | 3.164 | 0.0044 |

Table S4. Permutational multivariate analysis of variance (PERMANOVA) examining differences in sediment grain-size composition among reef positions and locations, including their interaction, based on Euclidean distances calculated from raw sediment grain-size proportions.

| Source | df | SS | MS | Pseudo-F | P(perm) |
|---|---|---|---|---|---|
| Position | 3 | 1.997 | 0.66565 | 58.182 | <0.0001 |
| Location | 2 | 0.74904 | 0.37452 | 32.735 | <0.0001 |
| Position × Location | 6 | 1.0562 | 0.17604 | 15.387 | <0.0001 |
| Site (Position × Location) | 0 | 0 | — | No test | — |
| Residual | 108 | 1.2356 | 0.011441 | — | — |
| Total | 119 | 5.0378 | — | — | — |

Table S5. Pairwise comparisons of sediment grain-size composition among reef positions within each location, based on Euclidean distances and permutation tests.

| Location | Comparison | Pseudo-t | P(perm) |
|---|---|---|---|
| Pompano Beach | Nearshore - Inner reef | 3.783 | 0.0016 |
| | Nearshore - Middle reef | 4.7871 | <0.0001 |
| | Nearshore - Outer reef | 6.5618 | <0.0001 |
| | Inner reef - Middle reef | 7.0126 | <0.0001 |
| | Inner reef - Outer reef | 9.0168 | <0.0001 |
| | Middle reef - Outer reef | 3.4346 | <0.0001 |
| Lauderdale Beach | Nearshore - Inner reef | 5.2532 | <0.0001 |
| | Nearshore - Middle reef | 3.0604 | 0.0005 |
| | Nearshore - Outer reef | 4.6922 | <0.0001 |
| | Inner reef - Middle reef | 6.2105 | <0.0001 |
| | Inner reef - Outer reef | 8.2251 | <0.0001 |
| | Middle reef - Outer reef | 2.5972 | 0.0009 |
| Hollywood | Nearshore - Inner reef | 2.4862 | 0.0006 |
| | Nearshore - Middle reef | 5.8804 | <0.0001 |
| | Nearshore - Outer reef | 5.2338 | <0.0001 |
| | Inner reef - Middle reef | 7.7309 | <0.0001 |
| | Inner reef - Outer reef | 8.0772 | <0.0001 |
| | Middle reef - Outer reef | 4.5236 | <0.0001 |

Table S6. Permutational analysis of multivariate dispersion (PERMDISP) examining differences in sediment grain-size variability among reef positions and locations.

| Analysis | df1 | df2 | F | P(perm) |
|---|---|---|---|---|
| Position | 3 | 116 | 41.186 | <0.0001 |
| Location | 2 | 117 | 0.32648 | 0.7332 |
| Pompano Beach - position | 3 | 36 | 2.1944 | 0.144 |
| Lauderdale Beach - position | 3 | 36 | 1.1944 | 0.3894 |
| Hollywood - position | 3 | 36 | 2.8743 | 0.0634 |
